# Altered Brain Extracellular Matrix Structure and Sex-Specific Learning/Memory Dysfunction in Mice Lacking the Functional Amyloid Cystatin-Related Epididymal Spermatogenic (CRES)

**DOI:** 10.64898/2026.05.27.728219

**Authors:** Alejandra Gomez, Aric F. Logsdon, Jeremy D. Bailoo, Gail A. Cornwall

**Author notes:** Correspondence should be addressed to: Gail A. Cornwall, Ph.D., Department of Cell Biology and Biochemistry, Texas Tech University Health Sciences Center, Lubbock, TX 79430. AG, AFL, JDB, GAC designed and performed research, analyzed data, wrote the manuscript. AFL, JDB, GAC acquired the funding.

## Abstract

The brain’s extracellular matrix (ECM) is highly specialized and unique. Unlike the collagen-rich matrices found in other tissues, it consists of complex proteoglycans and glycosaminoglycans that play critical roles in neuroplasticity including the integration and storage of new memories. Cystatin-related epididymal spermatogenic (CRES) is a member of a reproductive subgroup of family 2 cystatins and part of a specialized functional amyloid-containing ECM with host defense functions in the epididymal lumen. CRES and the endogenous epididymal amyloid matrix exhibit high plasticity, forming distinct amyloid structures, with antimicrobial functions tailored to specific bacterial strains. Recent studies show that CRES/CRES amyloid is also present in the mammalian brain ECM and suggest its inherent shape-shifting properties might contribute to neuroplasticity. In this study, we utilized a CRES knockout (KO) mouse model to determine if the loss of CRES altered the structure of the hippocampal and cortical ECM and impacted brain function. We report that CRES KO male and female mice exhibited sex-specific structural changes in both the loose/interstitial and perineuronal net (PNN) ECM. Additionally, young CRES KO males—but not females—displayed deficits in learning, memory and executive functioning when compared to WT controls. In contrast, older CRES KO males did not exhibit behavioral deficits, a finding that correlated with the observation of a similar PNN ECM structure when compared to WT mice. Our data suggest that CRES functions as a regulatory or structural component of the brain ECM, contributing to its sex-specific structural organization and plasticity, affecting sex-specific behaviors of learning and memory.

**Significance statement:** Structural changes in the brain ECM drive neuroplasticity that is essential for normal function; however, these same alterations can contribute to neurodegeneration and neurological disorders. Brain ECM structure differs between males and females and promotes sex-specific behaviors; however, the full extent of sex-specific distinctions in ECM structure/function relationships has yet to be elucidated. Using a CRES KO mouse model, we show that the functional amyloid CRES plays a regulatory/structural role in the brain ECM. The loss of CRES resulted in sex differences in loose/interstitial and PNN ECM organization and a male but not female learning and memory deficit. Our studies suggest that CRES/CRES amyloid contributes to ECM neuroplasticity and remodeling processes that drive sex-specific behaviors and possibly sexually dimorphic disease.

## Introduction

The brain extracellular matrix (ECM) is involved in critical processes across development by shaping synaptic plasticity and stability, cognitive flexibility, and influencing the overall physiology of the central nervous system (CNS) (Soles et al. 2023). The brain ECM is divided into several distinct populations including: 1) loose/interstitial ECM that surrounds neurons and glial cells; 2) membrane-associated ECM that surrounds blood vessels; and 3) highly specialized and condensed ECM labeled as perineuronal nets (PNNs) that surround distinct classes of fast firing excitatory and inhibitory neurons. The brain ECM plays key roles in neuroplasticity with the flexible loose/interstitial ECM allowing for rapid synaptic changes compared to the more restrictive PNNs. Although stable structures, the PNNs have become a particular focus of study because of their critical roles in regulating neuroplasticity; they act as a molecular “brake” that permits subtle, controlled remodeling necessary to integrate new memories without compromising the stability of existing neural circuits (Bozzelli et al. 2018; Sanchez et al. 2024; Fawcett et al. 2022). Changes in PNN structure, including its major structural component chondroitin sulfate proteoglycans (CSPGs) and the sulfation and glycan content of associated chondroitin sulfate glycosaminoglycans (CS-GAGs), are integral for PNN remodeling for neural functioning but are also associated with neurological disorders and neurodegeneration (Miyata and Kitagawa 2016; Scarlett, Hu, and Alonge 2022). While the mechanisms that underlie ECM remodeling remain unclear, the proteolytic breakdown and synthesis of new ECM molecules and recycling of its components are thought to contribute to ECM plasticity (Burk et al. 2022).

Although studies are still limited in number, there is evidence that the brain ECM is structurally different between males and females including in its mechanical properties and composition of collagens and other ECM proteins (Batzdorf et al. 2022). Indeed, the PNN structure, composition, and distribution, can vary between males and females in the hippocampus, cortex and amygdala influencing plasticity and resulting in sex-specific behaviors. Defining brain ECM structure/function relationships and if and how they differ between males and females is therefore key to elucidating mechanisms of sexually dimorphic behaviors and disease.

CRES is a member of a reproductive subgroup of family 2 cystatins of cysteine protease inhibitors that lack consensus sites for inhibition of papain-like cysteine proteases, suggesting distinctive functions (Cornwall and Hsia 2003; Cornwall, Orgebincrist, and Hann 1992). Subgroup members CRES, CRES2, CRES3 and cystatin E2 are highly amyloidogenic *in vitro* and contribute to the assembly of an amyloid-rich ECM that functions as a host defense structure to protect the male germline in the mouse epididymal lumen (Myers, Hastert, and Cornwall 2022; Whelly et al. 2012; Whelly et al. 2016). CRES and the epididymal amyloid matrix have shape-shifting properties, assembling into diverse amyloid structures with different host defense functions (e.g., bacterial trapping versus killing) depending on the bacterial strain (Myers, Hastert, and Cornwall 2022). Recently, it was shown that CRES and CRES amyloids are also part of the brain ECM including the loose/interstitial ECM and the condensed PNNs. Further, studies using a CRES knockout (KO) mouse model, showed that the loose/interstitial ECM from the male, but not the female hippocampus had reduced amyloid content compared to wildtype (WT) mice (Gomez et al. 2026). These data imply that the central roles played by CRES/CRES amyloids in the brain ECM varies between sexes.

Here, we evaluated whether the loss of CRES caused sex-specific alterations in ECM structure and brain function, with a focus on the hippocampus and cortex, using the CRES (KO) mouse model. Analysis of the morphology of the loose/interstitial ECM by transmission electron microscopy and of the condensed PNN ECM by wisteria floribunda agglutinin (WFA) and aggrecan staining showed that the ECM was structurally altered in a region-dependent and sex-specific manner in CRES KO mice compared to their WT counterparts. Further, the changes in structure correlated with a sex-specific behavioral phenotype, where young (18-25 wks) male but not young female CRES KO mice exhibited a learning and behavioral flexibility deficit. In contrast, older (60-67wks) male KO mice did not exhibit the behavioral phenotype and PNN structure resembled that of WT animals suggesting a recovery of PNN structure/function with age. Together, our data showcase CRES/CRES amyloid is an important regulatory or structural component of the brain ECM that contributes to sex-specific behaviors and possibly sexually dimorphic disease.

## Materials and Methods

### Animals

All animal studies were conducted in accordance with the Guide for the Care and Use of Experimental Animals under a protocol approved by the University Institutional Animal Care and Use Committee. Female and male 129SvEv/B6 WT and CRES knockout mice were bred in-house. All mice were group-housed and maintained under a constant 12-hour light/12-hour dark cycle with food and water available *ad libitum*. Nestlets and plastic tunnels were provided to the animals as additional cage resource supplementation. Female mice weighed between 19–22 g at 18–24 weeks and 24–34 g at 60–67 weeks. Male mice weighed between 17–27 g at 18–24 weeks and between 32–40 g at 60–67 weeks of age. A total of 80 mice were used for behavioral testing and 36 mice for extracellular matrix analyses. Mice were euthanized by CO_2_ (3lpm) followed by cervical dislocation. Brains were separated into hemispheres sagittally, and either cryopreserved or dissected into regions (hippocampus and cortex) for ECM enrichment.

### Enrichment of brain ECM Populations

Biochemical enrichment of the different populations of hippocampal and cortical ECM including loose/interstitial (E1), membrane-associated (E2), ionic bound membrane-associated (E3), and perineuronal net (E4) was carried out using a protocol modified from that of Deepa et al., (Deepa et al. 2006). In the current report only the loose (E1) and PNN (E4) fractions were studied. Briefly, the dissected mouse brain regions from two mice were placed into E1 buffer (50 mM Tris pH 7.4, 150 mM NaCl, 2mM EDTA pH 8, 2 mM PMSF) (0.5-2 ml depending on brain region) on ice. The tissues were gently homogenized by passing sample through a 20G needle (∼10-15 pulls). The samples were centrifuged at 20,000 x g 30 min at 4°C. The supernatant was removed, the tissue pellet was extracted again in E1 buffer and the resulting supernatant was pooled with the first to generate the E1 fraction containing loose/interstitial ECM. The pellet was resuspended in E2 buffer (Buffer 1 plus 0.5% Triton X-100), centrifuged at 20,000 x g for 30 minutes at 4°C and extracted a second time to generate the E2 fraction containing membrane-associated ECM (E2). The resulting pellet was resuspended in E3 buffer (50 mM Tris pH 7.4, 1 M NaCl, 2 mM EDTA pH 8, 0.5% Triton X-100, 2 mM PMSF), centrifuged at 20,000 x g for 30 minutes at 4°C and the resulting supernatant contained ionic bound membrane-associated ECM (E3). The remaining pellet was resuspended in E4 buffer (Buffer 2 plus 7M urea), centrifuged at 20,000 x g for 30 minutes at 4°C and the supernatant contained the PNN-tightly bound ECM-(E4) fraction. The protein concentration of each extraction was determined by BCA assay (Sigma/Millipore, St. Louis, MO).

### Preparation of loose/interstitial ECM insoluble extract

One milligram of loose/interstitial (E1) extract from cortex and hippocampus from WT and CRES KO male and female mice was centrifuged at 225000 x g for 70 min at 4°C using a tabletop Beckman Optima TLX ultracentrifuge to generate the supernatant (S) and pellet containing insoluble material (E1 pellet). The resulting pellets were resuspended in E1 buffer and used for Transmission Electron Microscopy (TEM). Protein concentrations were determined by BCA assay (ThermoScientific, Rockford).

### Negative Stain TEM

Three μgs of hippocampal E1 pellets were spotted onto Formvar/carbon-coated 200-mesh nickel grids (Ted Pella, Redding, CA) for 5 min. The samples were wicked off with filter paper and washed for 1 minute with water, stained with 2% uranyl acetate for 1 minute, washed twice with water for 1 minute, and then dried for 2 minutes. Grids were examined with a Hitachi 7650 electron microscope. For image quantification analysis a minimum of four representative TEM micrographs from male and female wild-type and CRES knockout E1 hippocampal extracts (E1 total and E1 pellets) from three independent experiments were imported as TIFF files into ImageJ and converted to 8-bit images. Individual micrographs were quantified to identify the percentage area of the micrograph that contained structure. A threshold was applied across all images to remove background from structures of interest. Outlines of structures were generated using the Selection tool and used to measure area and relative intensity. Finally, the percentage area was calculated by measuring the area of the selection relative to the total image area. Values for individual images were averaged per experiment. Two-way ANOVA were performed using GraphPad/Prism version 10.

### PNN staining

Brain hemispheres from 129SvEv/B6 female and male (18-24 and 60-67 weeks old) mice were cryopreserved in Cryomatrix (Epreda Kalamazoo, MI). Fifteen µm sagittal sections were mounted on to Superfrost Plus microscope slides (Fisher Scientific, Pittsburg PA) and stored at -80°C until use. The slides were thawed at room temperature for 30 minutes in a Tupperware container containing damp filter paper. Tissue sections were outlined with a PAP pen and rehydrated with 1x Dulbecco’s phosphate buffered saline (DPBS) (Sigma-Aldrich St. Louis MO, without calcium chloride) for 10 minutes and then fixed with 4% methanol-free formaldehyde (Thermo Scientific, Rockford IL, # 28908) in DPBS for 1 hour, at room temperature. Sections were washed with 1x DPBS three times for 2 minutes each and permeabilized with 0.2% Triton X-100 (ultrapure, Thermo Scientific, Rockford IL, cat # 28314) in DPBS for 5 minutes. Sections were washed with DPBS then blocked with 30% heat inactivated goat serum (normal goat serum, Vector Labs, Newark CA, cat # S-1000, prepared by incubating goat serum at 56°C for 45 min followed by filtration through a 2 µm filter) in DPBS for 1 hour. Serum was clarified before each use by centrifugation at 15,700 x g for 10 min at 4°C. Sections were incubated with anti-Aggrecan rabbit polyclonal antibodies (AB1031 CHEMICON Merck KGaA, Darmstadt Germany), 5 µg/ml in 30% heat inactivated goat serum in DPBS), overnight, at 4°C. After, sections were washed with DPBS three times, 2 minutes each, then incubated with Alex Fluor 594-labeled anti-rabbit secondary antibody (Invitrogen, Waltham, MA), diluted 1:500 in 30% heat inactivated goat serum in DPBS for two hours at room temperature in the dark. Sections were washed with DPBS then incubated with 0.5 µM DAPI (Molecular Probes, Invitrogen Eugene, OR) in DPBS for 10 minutes. Finally, sections were washed with DPBS three times for 2 minutes each followed by a wash with deionized water for 2 minutes. Sections were mounted with one drop of VectaMount Aqueous mounting medium (VectorLabs, Newark, CA) and a coverslip (22×22 mm, No. 1.5) was used to cover the tissue. The excess mounting medium and any air bubbles were removed by gently pressing on the coverslip. Subsequently, the sections were dried for 30 minutes and sealed with nail polish. For *Wisteria floribunda* agglutinin (WFA) staining, cryosections slides were fixed, permeabilized and blocked as described previously, then incubated with FITC-conjugated WFA (cat #32481 Life Technologies, Eugene, OR) diluted 1:200 in 30% heat inactivated goat serum for 2 hours at room temperature in the dark. Sections were washed with DBPS three times 2 minutes each, counterstained with DAPI and mounted as described above.

Stained perineuronal nets were imaged from cortex and hippocampus of sagittal mouse brain sections. The cerebral cortex was divided into four main regions: frontal, cingulate, parietal, and occipital as indicated in Fig SI3. Images were obtained using a Zeiss 200 Axiovert 200M fluorescent microscope with Zen Blue version 3.5. All images were acquired using the same parameters.

### Fluorescence intensity quantification

The relative fluorescence intensity (RFU) levels of WFA- and aggrecan-stained cortex and hippocampus tissue sections from 18-to 24-week-old and 60- to 67-week-old male and female mice were quantified using ImageJ. Single-channel images were imported into FIJI, converted to 8-bit, and a threshold of 20/80 was applied to minimize background signal. The same threshold values were used within the same experiment for comparisons between ages and sex. Individual PNNs were selected as regions of interest, and pixel values of PNNs were measured in brain tissue from six females and six males per age group (3 WT, 3 CRES KO), which were then averaged/experiments for a total of 3 experiments. The total number of WFA- or ACAN-positive PNNs was determined by manual counting. Statistical analyses were performed using GraphPad Prism Version 10. Each age group was analyzed separately using a two-way ANOVA with genotype and sex as fixed effects.

### CS-GAG sulfation pattern analysis

The E1 pellets from cortex and hippocampus were generated as described above. All pellets were resuspended in sterile water and centrifuged a second time, under the same conditions, to ensure that all salt was removed. The resulting pellet was resuspended in 50 µl of sterile water. The E4 fractions were dialyzed against water with three changes to remove salt using Slide-A-Lyzer mini dialysis units (cat #ZD389330 Thermo Scientific, Rockford IL). Protein concentrations were determined by BCA assay (ThermoScientific, Rockford, IL). Fifty µg of sample were digested in a rotating incubator set to 80 rpm at 37°C for 24 h using 500 mU/mL Chondroitinase ABC, (ChABC cat #C3667, Sigma St Louis, MO.) reconstituted in 50 mM ammonium bicarbonate (pH 7.4). Enzyme digested samples (100µl) were added to a 3kDa MWCO Amicon spin-columns (Sigma) with 100µl LC-grade optima water (Fisher Scientific), and centrifuged for 10 min at 14,000 × g. The flow-through was collected and dehydrated using a Vacufuge Plus (Eppendorf), and the dried product was reconstituted in 40 μL of optima water.

CS-GAG sulfation patterns were measured using liquid chromatography tandem mass spectrometry (LC-MSMS) methods (Alonge et al. 2019). Briefly, 5 μL of CS-GAG disaccharides were resolved through a 50°C Hypercarb column (2.1 × 50 mm, 3 μm, Thermo Fisher) at 0.3 mL/min using an Exion AE LC-40 paired with a triple quadrupole mass spectrometer QTRAP 6500+ (AB Sciex) equipped with an electrospray ion source (negative mode) and operated in multiple reaction monitoring (MRM) unit mass mode using the following conditions: the electrospray voltage was set to −4.5 kV, the curtain gas was set to 25 psi, the CAD gas was set to 9 psi, and the source temperature was 450°C. Precursor and fragment transitions and collision energies (CE ranging from 5 to 40) for each MRM channel were optimized and found to be Δ4S-CS (CS-A) *m/z* 458 > 300 using a CE of -30, Δ0S-CS (CS-O) *m/z* 378 > 175 using a CE of -16, and Δ6S-CS (CS-C) *m/z* 458 > 282 using a CE of -27. The declustering potential (DP) of both Δ4S-CS (CS-A) and Δ6S-CS (CS-C) was set at -60, and -30 for Δ0S-CS (CS-O). CSPG isomer separation was achieved using the following buffers: Buffer A = 100 mM ammonium acetate, pH 9.5 optima water, buffer B = 0.1% ammonium hydroxide in LC-grade optima acetonitrile (Fisher Scientific), and the following gradient: 100% A held for 3 min, then increased to 5% B over 10.5 min. B was then increased from 5% to 90% from 10.5 to 11.5 minutes. The column was then washed with 90% B for 1 min (11.5-12.5 min) and re-equilibrated at 100% A for 3 min, for a total run time of 15.5 min. This was found to provide optimal chromatographic resolution for each CS-GAG isomer.

Industry-grade CS-GAG disaccharide standards Δ0S-(OC28897), Δ4S-(OC28898) and Δ6S-(OC01702) were purchased from Biosynth (Berkshire, UK). SciexOS software was used to acquire and quantify the ratios between peak areas consistently produced from equimolar CS-GAG standard runs throughout the detectable range of quantification when normalized to the highest peak intensity. Relative quantification of each CS-GAG isomer within a sample was achieved using a modified peak area normalization function (Alonge et al. 2019).

### Behavioral evaluations

Young (18–24 weeks) and old (60–67 weeks) mice were tested for differences in learning and memory (simple discrimination) and executive functioning (reversal learning) using a novel, validated water T-maze protocol (Bailoo et al. 2024). Briefly, animals underwent a water-based, two-choice discrimination task across three days (six trials per day). First, animals learned to discriminate the location of a platform in one of two possible arms using spatial cues across nine trials (simple discrimination). On day 2, trial 4, the platform was switched to the opposite arm (reversal learning) and remained there for subsequent trials. Our primary outcome variables were latency to the platform and the cumulative number of incorrect arm entries per stage.

Because age-related motor impairment is common in mice, we also evaluated motor ability in a subset of animals (n = 5 per sex, genotype, and age) to rule out motor deficits as a confounding factor for differences between CRES KO and WT mice. Animals completed a voluntary exploration task (open-field test) and a forced motor coordination task (accelerating rotarod) using established protocols (Bohlen et al. 2014; Spontarelli et al. 2024). Primary outcome variables were distance traveled in the open field, and latency to fall off the accelerating rotarod. Mice were euthanized 7–10 days post-behavioral testing for extracellular matrix enrichment or brain cryopreservation, as previously described.

Statistical analyses were performed using IBM SPSS version 31. For all behavioral evaluations, young and old cohorts were analyzed separately. In the water T-maze, stages (simple discrimination, reversal learning) were also analyzed separately.

We modeled latency to the escape platform using a linear mixed-effects model. Trial was nested within mouse using an Autoregressive 1 (AR1) covariance structure. A restricted maximum likelihood (REML) estimation procedure was used, with a Satterthwaite approximation and a maximum of 100 iterations (10 maximum step-halvings). Genotype and sex were included as fixed effects, and significant effects were probed using sequential Bonferroni-corrected pairwise comparisons.

To directly distinguish whether increases in latency to reach the escape platform reflect true deficits in spatial learning rather than decision hesitancy or altered motor output, choice accuracy in the water T-maze was additionally evaluated. The cumulative number of incorrect arm entries was calculated across sequential trial blocks (3 trials per block) to generate a trial-block learning curve. Choice accuracy data were analyzed using a three-way repeated measures analysis of variance (ANOVA). Block (Blocks 1, 2, and 3) was entered as the within-subjects repeated measure, with Genotype (WT, CRES KO) and Sex (male, female) included as between-subjects factors. The assumption of sphericity was assessed via Mauchly’s test; where this assumption was violated, degrees of freedom were adjusted using the Greenhouse-Geisser correction. Significant interactions, including the Block × Genotype × Sex interaction, were probed using simple main effects and Bonferroni-corrected pairwise comparisons to determine if spatial learning trajectories diverged between sexes.

For analysis of the open-field and rotarod data, two-way ANOVAs with genotype (WT, CRES KO) and sex (male, female) were used. Significant main effects or interactions were probed with Bonferroni-corrected post hoc tests. Model assumptions were examined following established methods (Bohlen et al. 2014; Spontarelli et al. 2024; Bailoo et al. 2024).

## Results

### Sex-specific differences in loose/interstitial hippocampal ECM morphology in CRES KO mice

Studies were carried out to determine if the morphology of the loose/interstitial ECM changed with the loss of CRES. Using an established protocol to enrich for different populations of brain ECM (Deepa et al. 2006), the loose/interstitial ECM (E1 extract) was isolated from young (∼18-25 wks) CRES wildtype (WT) and CRES KO male and female hippocampus and separated into its soluble (supernatant) and insoluble (E1 pellet) components by centrifugation. The insoluble pellet was then examined by negative-stain transmission electron microscopy (TEM). The hippocampal E1 pellet has been shown to be enriched in thioflavin T positive material indicating the presence of cross β structure typical of amyloid (Gomez et al. 2026).

The hippocampal E1 pellet from male and female WT mice was characterized by large clusters of a highly branched matrix (Fig 1) that contained small ball-like structures (Fig SI1, pink arrows). Fibrillar and film-like structures were also occasionally present (Fig SI1, blue arrows). In contrast to the visually comparable structures observed in the male and female WT mice, a sex-specific difference in ECM morphology was observed in the E1 pellets from CRES KO mice (Fig 1, Fig SI1). CRES KO males displayed small matrix fragments (Fig 1) which at higher magnification (20,000X) revealed structures characteristic of protofibrils and oligomers, suggesting an immature/incomplete assembly of the branched matrix (Fig SI1, green arrows). The smaller matrix fragments could also indicate an ECM that partially disassembled or fragmented during biochemical isolation. While some small matrix fragments were also detected in the CRES KO female, large, disorganized tangles/masses containing ball-like structures were common, suggesting an abnormal ECM structure, but with properties different from that in the male, possibly indicative of an aberrant aggregation of the ECM components (Fig 1, Fig SI1). Further, the ball-like structures associated with the disorganized masses were often large in the KO females. Similar sex-specific differences in morphology were observed in loose/interstitial hippocampal ECM that had not been centrifuged (total E1 ECM) indicating the distinct structures were due to genotypic and sex differences and not a result of the concentration of the insoluble material during centrifugation (Fig SI2). Male CRES KO mice displayed small matrix fragments while female CRES KO mice showed larger assemblies including fibrillar bundles and tangles of matrix/balls (Fig SI2). To quantify the changes in ECM between CRES WT and KO mice, the area occupied by structure relative to the total area of the TEM micrograph and the relative intensities of the individual structures were determined using ImageJ. As shown in Fig 1B, the area occupied by the branched matrix in WT males was significantly larger compared to that occupied by the smaller and dispersed matrix fragments in KO males (*p =* 0.032). While the area occupied by the large masses in the female KO group was not different from that occupied by the branched matrix in the WT females, the masses exhibited significantly higher intensity compared to WT females (*p =* 0.007). Together our results show that CRES is important for normal hippocampal ECM structure and suggest that the physiochemical and mechanical properties of the loose/interstitial hippocampal ECM and the contribution of CRES to its structure varies between sex.

**Figure 1.**
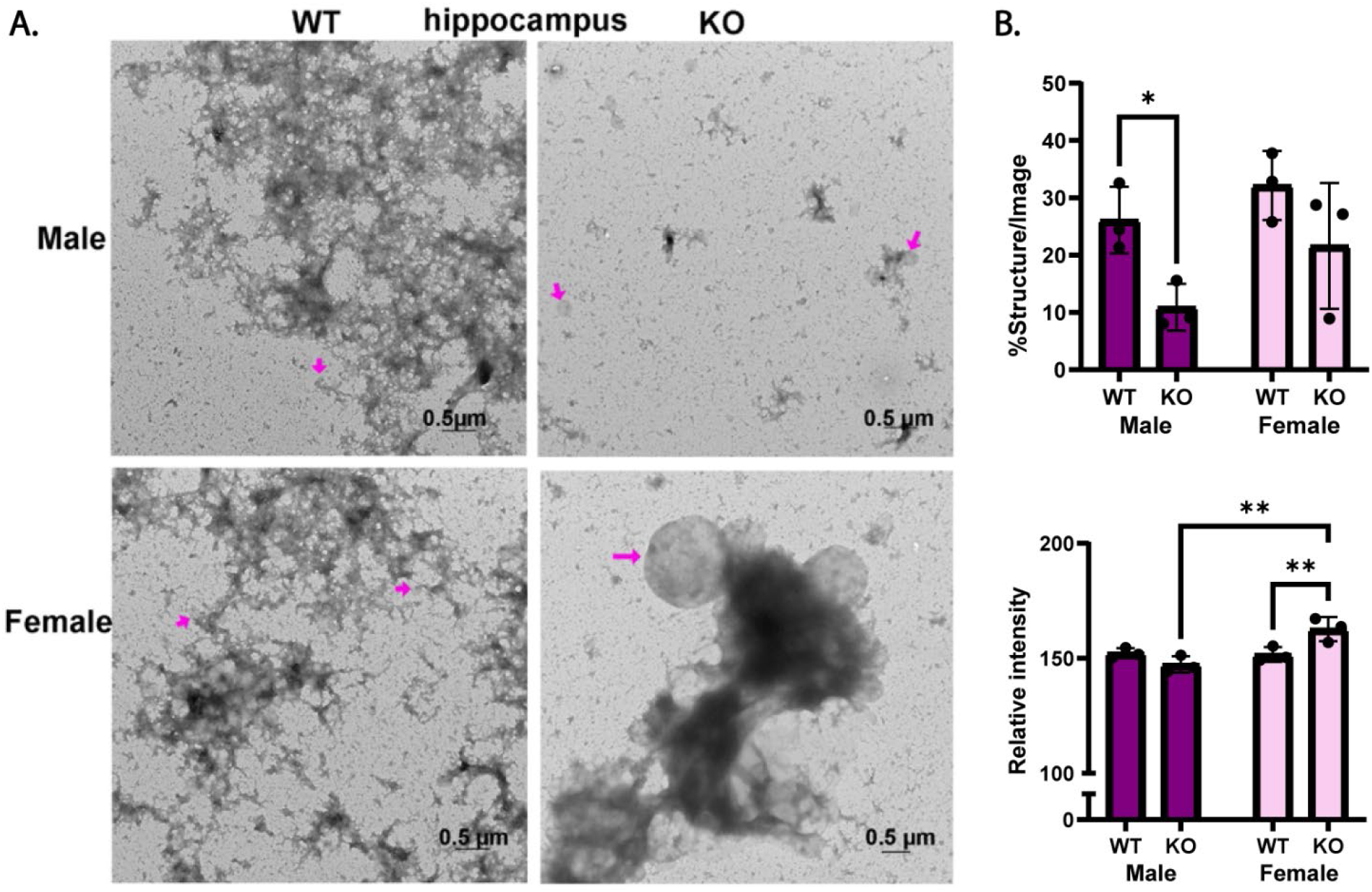
Sex-specific differences in loose/interstitial hippocampal ECM morphology in CRES KO mice. **A)** Negative stain transmission electron microscopy of the loose/interstitial ECM pellet (E1) showed that a branched matrix with small associated ball-like structures (pink arrows) was the predominant structure in extracts from male and female CRES WT mice. In contrast, CRES KO males showed small fragments while KO females exhibited small matrix fragments and large masses with associated, and often large, ball-like assemblies, possibly representing ECM aggregates. Scale bar, 0.5 µm. n=3-4 independent experiments using E1 pellets isolated from 3-4 different hippocampal ECM preparations. Additional images are in Fig S1 and S2. **B)** Quantification of ECM structure area and intensity. At least four TEM micrographs of both E1 total and E1 pellet extracts per mouse group were quantified from 3-4 independent experiments. Analysis of variance showed that the area occupied by the branched matrix in WT males was significantly larger than that observed in CRES KO males (*p* = 0.032). Although no significant difference in area was detected between WT and KO females, ECM structures in the KO group exhibited significantly higher intensity compared to WT females (*p* = 0.007) and KO males (*p* = 0.001), consistent with the presence of large masses.

### PNN glycan and protein components are altered in young CRES KO mice

CRES is expressed by PNN-containing neurons and is found in brain extracts enriched in PNN ECM proteins (Gomez et al. 2026); therefore, we next asked whether PNNs were compromised in the absence of CRES. Using brains from WT and CRES KO males and females at a young (20-24 weeks) and old (60-64 weeks) age we examined hippocampal and cortical PNNs through fluorescence microscopy using two strategies: 1) wisteria floribunda agglutinin (WFA), a lectin that binds to GalNAc β 1-3 Gal residues found along the disaccharide chains of CS-GAGs, thus looking at the glycan component of PNNs, and 2) anti-aggrecan (ACAN) 1031 monoclonal antibody that binds to the CSPG protein core of ACAN that make up PNNs, thereby looking at their most abundant protein component. Fig SI3 presents a schematic of the general division we adopted for the cerebral cortex. Along the sagittal plane, we examined PNNs originating from the frontal, cingulate, parietal, and occipital cortices, as well as the hippocampus and recorded both the relative fluorescence intensity (RFU) for WFA and ACAN, as well as the number of WFA positive (WFA^+^) and aggrecan positive (ACAN^+^) PNNs.

We measured significantly reduced levels of WFA RFU in the hippocampus and cortical regions of young male and female CRES KO animals compared to WT suggesting CRES contributes to the structure, and possible function of PNNs, throughout the cortex and hippocampus (Fig 2A, B). The number of WFA+ PNNs, however, did not differ between WT and KO mice except in the KO male hippocampus where fewer WFA positive PNNs were detected compared to WT males and KO females (Fig 2B). Although ACAN staining was measured independently from that of WFA, and therefore does not represent the same PNN, no differences were observed in the ACAN RFU and number of ACAN+ PNNs between WT and CRES KO hippocampus and cortical regions, indicating PNNs were present and that CRES has a greater impact on the glycan composition of the PNN structure (Fig 2C). An exception to this was in the male CRES KO hippocampus where fewer ACAN+ PNNs were present compared to WT males and KO females, suggesting a more profound effect of the loss of CRES on PNN structure in the male hippocampus compared to females (Fig 2C). A sex-difference was also observed in the parietal and occipital cortex where WT females possessed significantly more ACAN positive PNNs compared to WT males (Fig 2C). This sex difference was lost in KO animals because of a profound drop in the number of ACAN positive PNNs in these two regions in the KO females resulting in a number of ACAN positive PNNs like that in KO males (parietal cortex *p =* 0.0002, occipital cortex *p <* 0.0001 compared to WT). The change in the number of PNNs may reflect a loss of PNNs, an altered PNN structure possibly due to masking of aggrecan epitopes, and/or the presence of PNN populations that are aggrecan negative. Overall, our data suggest that in young mice PNN structure is altered in the absence of CRES, with some region-dependent sex differences and that the male hippocampus and discrete regions of the female cortex are especially susceptible.

**Figure 2.**
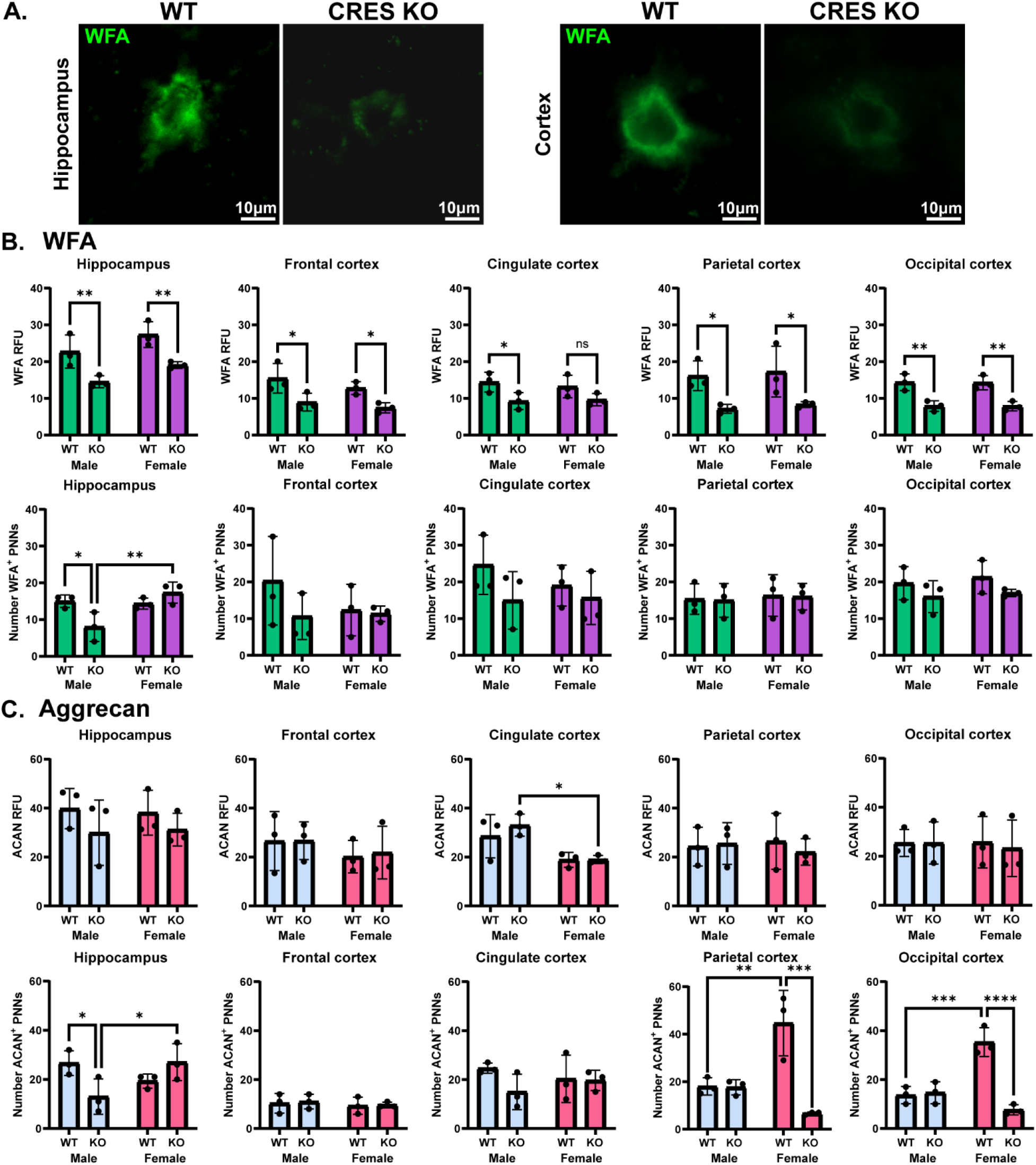
PNN glycan and protein components are altered in young CRES KO mice. **A)** WFA (green fluorescence) staining of hippocampal and cortical PNNs from female CRES WT and KO mice. Scale bar: 10 µm. **B)** WFA relative fluorescence intensity (RFU) and number of WFA+ PNNs in hippocampal and cortical PNNs from male (green bars) and female (purple bars) CRES WT and KO mice. The WFA RFU was significantly decreased in the hippocampus and all cortex regions from CRES KO male and female mice compared to WT (Male, hippocampus *p* = 0.009, frontal cortex *p* = 0.016, cingulate cortex *p* = 0.035, parietal cortex *p* = 0.027, occipital cortex *p* = 0.002; Female, hippocampus *p* = 0.009, frontal cortex *p* = 0.035, parietal cortex *p* = 0.027, occipital cortex *p* = 0.002). However, the number of WFA+ PNNs was significantly reduced only in the male hippocampus compared to WT male (*p* = 0.014) and KO female (*p* = 0.003). **C)** ACAN RFU and number of ACAN+ PNNs in hippocampal and cortical PNNs from male (blue bars) and female (red bars) CRES WT and KO mice. The ACAN RFU was not different between male and female WT and CRES KO mice except in the cingulate cortex where decreased intensity was observed in KO females compared to KO males (*p* = 0.012). The number of ACAN+ PNNs was significantly reduced in the male KO hippocampus compared to WT male (*p* = 0.023) and KO female (*p* = 0.021). A sex difference was observed in the parietal (*p* = 0.002) and occipital cortex (*p* < 0.001) where WT females possessed significantly more ACAN+ PNNs compared to WT males. This difference was lost from the KO females as they displayed significantly fewer ACAN+ PNNs (parietal, *p* < 0.001, occipital, *p* < 0.001) compared to WT females. Values represent Mean ± SD from n = 3 biological replicates.

### The loss of CRES does not affect CS-GAG Sulfation

Varying levels of CS-GAG sulfation are known to regulate PNN structure and function (Huang et al. 2023; Dick et al. 2013; Fawcett et al. 2022; Belliveau et al. 2024). Because we show a reduction in the intensity of WFA in PNNs in the brain tissue of CRES KO mice (Fig 2), we next used LC-MSMS to analyze the relative abundance of three CS-GAG isomers (CSO, CSC, CSA) in the loose/interstitial ECM E1 pellet and the condensed PNN fraction (E4 extract) from young male and female WT and CRES KO mice. The absence of CRES in the E1 pellet had no effect on the relative abundance of each of the three CS isomers in the cortex and hippocampus (Fig 3Ai) or on the total CS-GAG sulfation as indicated by the consistent levels of unsulfated CSO in the E1 pellets (Fig 3Aii). The absence of CRES also did not affect CS-GAG sulfation patterning, as indicated by a lack of change in the CSA:CSC ratios among the E1 pellets (Fig 3Aiii). Similarly, the absence of CRES in the hippocampal and cortical PNN ECM (E4 extract) had no effect on the relative abundance of each CS isomer measured (Fig 3Bi), total CS-GAG sulfation (Fig 3Bii), or on CS-GAG sulfation patterning (Fig 3Biii). In keeping with recent reports of brain region-specific differences in total CS-GAG sulfation (Belliveau et al. 2024), a two-way ANOVA revealed an overall effect of brain region [*F (3, 16)* = 5.705*; p =* 0.0075] on the relative abundance of unsulfated CSO levels among the hippocampal PNN fractions, compared to the cortical PNN fractions (Fig 3Aii). Of note, the effect of brain region on CSO levels was not observed in the loose/interstitial ECM E1 pellets (Fig 3Bii; [*F (3, 15) =* 2.080*; p =* 0.146]), suggesting that hippocampal PNNs (E4), but not hippocampal loose/interstitial ECM (E1), exhibit higher levels of CS-GAG sulfation compared to cortical PNNs, even in the absence of CRES. No sex differences were observed in CS-GAG sulfation patterns in relationship to a specific genotype, brain region or ECM fraction.

**Figure 3.**
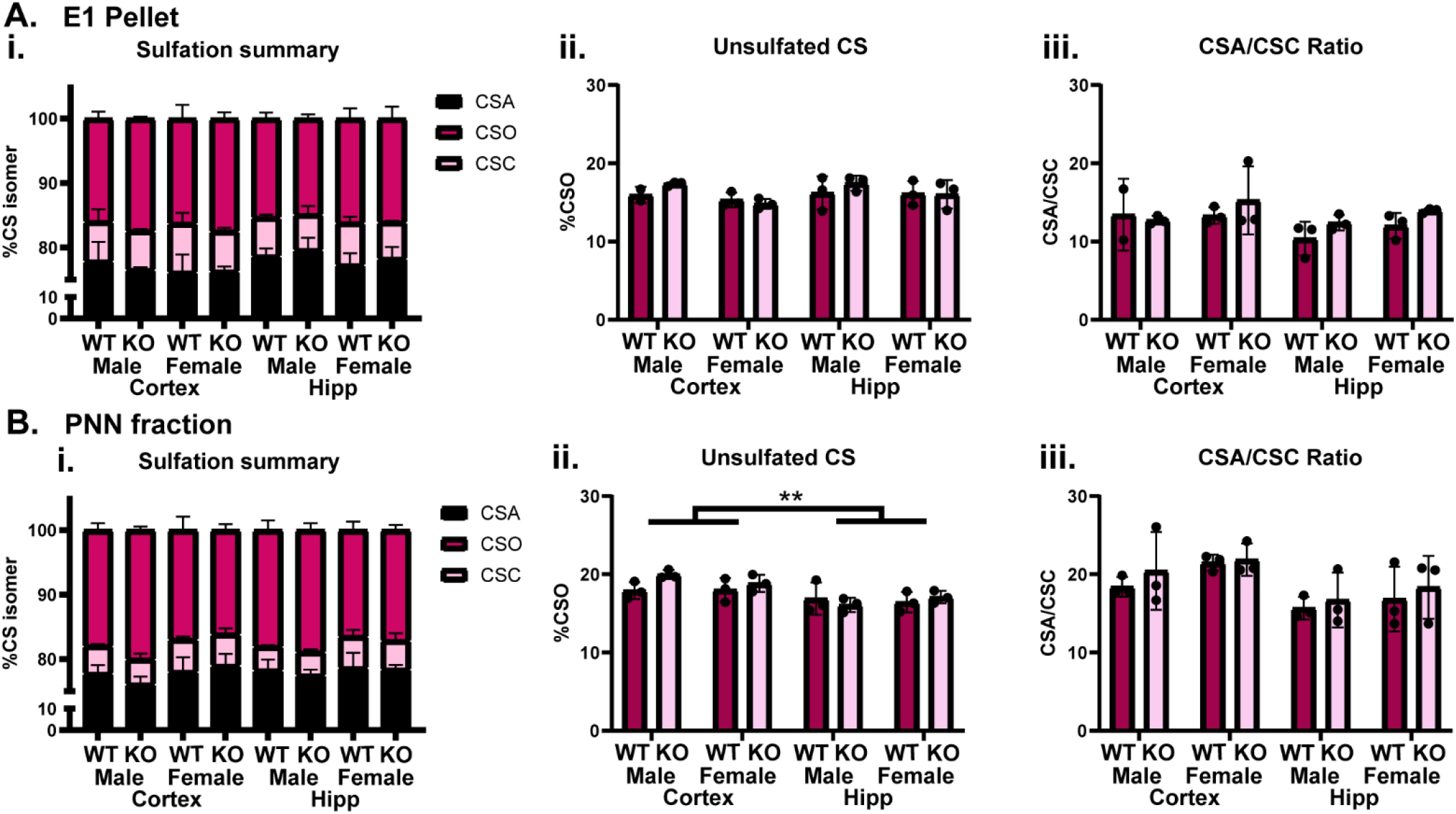
The loss of CRES does not affect CSPG sulfation in the loose/interstitial insoluble fraction (E1 pellet) or in the condensed PNN (E4) fraction. LC-MS/MS was used to measure sulfation in biochemical extracts of hippocampal (Hipp) and cortex: A) loose/interstitial E1 pellet and B) PNNs from male and female WT (dark pink) and CRES KO (light pink) young mice: i) Relative abundance of three CS-GAG isomers (CSO, CSC, CSA); ii, total CS-GAG sulfation as measured by relative abundance of unsulfated CS; and iii, CSA/CSC ratio indicating sulfation patterning. No genotype or sex differences were observed. However, a significant two-way ANOVA main effect of brain regions on total sulfation was observed in the PNN fraction where the hippocampus contained significantly more sulfated CS-GAGs (based on decreased unsulfated CS) compared to PNNs in the cortex [*F* (3, 16) = 5.705, *p* = 0.0075]. Values represent Mean ± SD of n = 3 biological replicates.

### Recovery of PNN structural changes in old CRES KO mice

In contrast to the alterations in WFA and ACAN observed in the young CRES KO mice, old KO mice showed fewer differences from WT. Indeed, the WFA RFU in old CRES KO mice matched that of WT mice throughout all brain regions (Fig 4A). The exception was in the hippocampus where the old KO males displayed fewer WFA+ PNNs compared to WT males (Fig 4B). Similarly, the ACAN RFU and number of ACAN+ PNNs were not different between old WT and KO animals with the exception of the KO male occipital cortex where a significant decrease in the number of ACAN+ PNNs were observed (Fig 4C, D). In addition, the number of ACAN+ PNNs was not different between old female KO and WT parietal and occipital cortices (Fig 4), a considerable change from that observed in the young KO females (Fig 4D). Our results suggest an age-related recovery of PNN structure in CRES KO mice. Alternatively, PNN development may be delayed in the younger CRES KO mice, but the aging process may usurp any CRES-related effects on adult PNN structural maturity.

**Figure 4.**
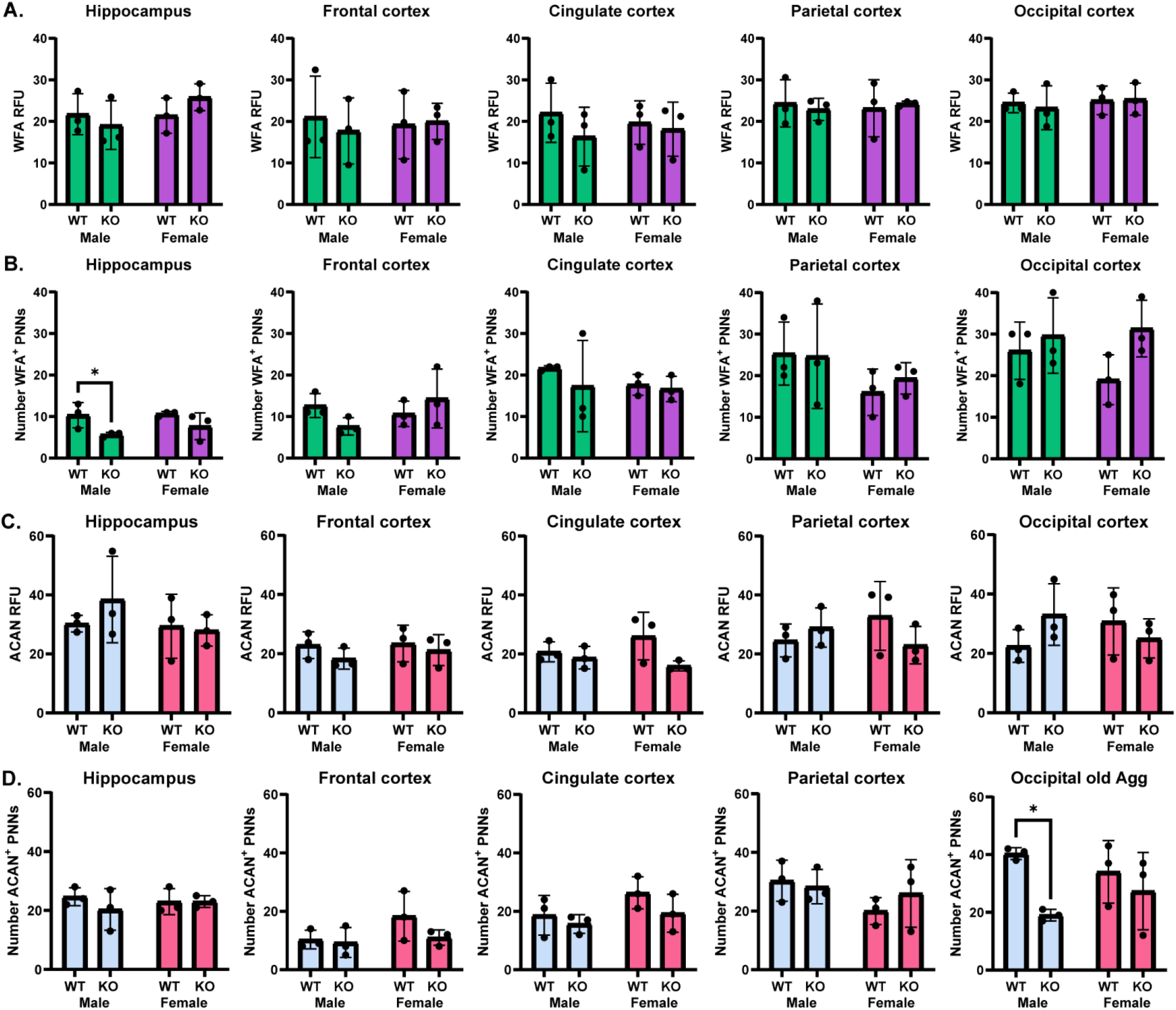
Recovery of PNN structure in old (60-64 wks) CRES KO mice. **A)** WFA relative fluorescence intensity (RFU) and **B)** number of WFA+ PNNs in hippocampal and cortical PNNs from old male (green bars) and female (purple bars) CRES WT and KO mice. No statistical differences in genotype were found except in the old male KO hippocampus which contained fewer WFA+ PNNs compared to WT males (*p* = 0.035). **C)** ACAN RFU and **D)** number of ACAN+ PNNs in hippocampal and cortical PNNs from old male (blue bars) and female (red bars) CRES WT and KO mice. No differences were found except in the old KO male occipital cortex where there was a decrease (*p* < 0.001) in the number of ACAN+ PNNs compared to WT males. Values represent Mean ± SD of n= 3 biological replicates.

We also observed that the overall WFA RFUs were often higher in the cortical PNNs of older WT mice (∼20-30 RFUs) compared to their younger counterparts (∼15-25 RFU) (Fig 5). An age-related effect on WFA levels has been described previously and is thought to represent an accumulation of PNN components (CSPGs) as a result of reduced microglial degradation or changes in glycan composition (Colon, Chan, and Maguire-Zeiss 2025). Despite these established age-related differences in WFA, in several regions the net increase in WFA intensity with age was often significantly greater in KO animals compared to that in WT suggesting compensatory mechanisms contributed to the recovery of PNN structure (Fig 5). Further these potential compensatory responses were more often observed in female as opposed to male KO mice suggesting sex-specific differences in PNN recovery mechanisms.

**Figure 5.**
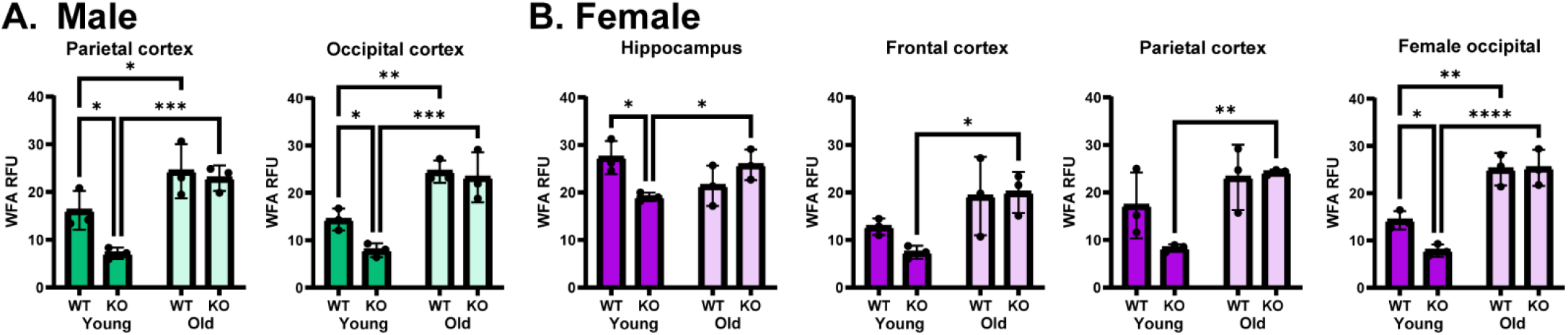
Compensatory responses in old CRES KO mice contribute to the recovery of PNN structure. PNNs from **A)** male and **B)** female brain regions in old CRES KO mice showed larger net increases in WFA RFU intensity than old WT mice when compared to their younger counterparts. Male parietal cortex: young vs old WT *p* = 0.028; young vs old KO *p* < 0.001; young WT vs KO *p* = 0.019. Male occipital cortex, young vs old WT *p* = 0.005; young vs old KO p < 0.001; young WT vs KO *p* = 0.038. Female hippocampus, young vs old KO *p* = 0.033; young WT vs KO *p* = 0.013. Female frontal cortex, young vs old KO *p* = 0.012. Female parietal cortex, young vs old KO *p* = 0.004. Female occipital cortex, young vs old WT *p* = 0.002; young vs old KO *p* < 0.001; young WT vs KO *p* = 0.024. Values represent Mean ± SD of n = 3 biological replicates.

### Young CRES KO males but not females show deficits in cognition and behavioral flexibility

Young and aged WT and CRES KO mice of both sexes (n = 10 per group) were evaluated using a novel validated water T-maze two-choice discrimination paradigm (Bailoo et al. 2024). In younger mice, compared to the WT controls, CRES KO males—but not females—displayed a deficit in learning and memory (simple discrimination), taking, on average, a significantly longer time (16.658 s) to learn to locate the escape platform [Genotype x Sex interaction: *F* (1, 45.408) = 4.245, *p* = 0.045]. A similar, yet more pronounced deficit in executive functioning during reversal learning was observed. When compared to WT mice, CRES KO males—but not females—took on average, a significantly longer time (24.416 s) to learn to locate the escape platform [Genotype x Sex interaction: (*F* (1, 33.294) = 4.908, *p* = 0.034)] (Fig 6A-B).

**Figure 6.**
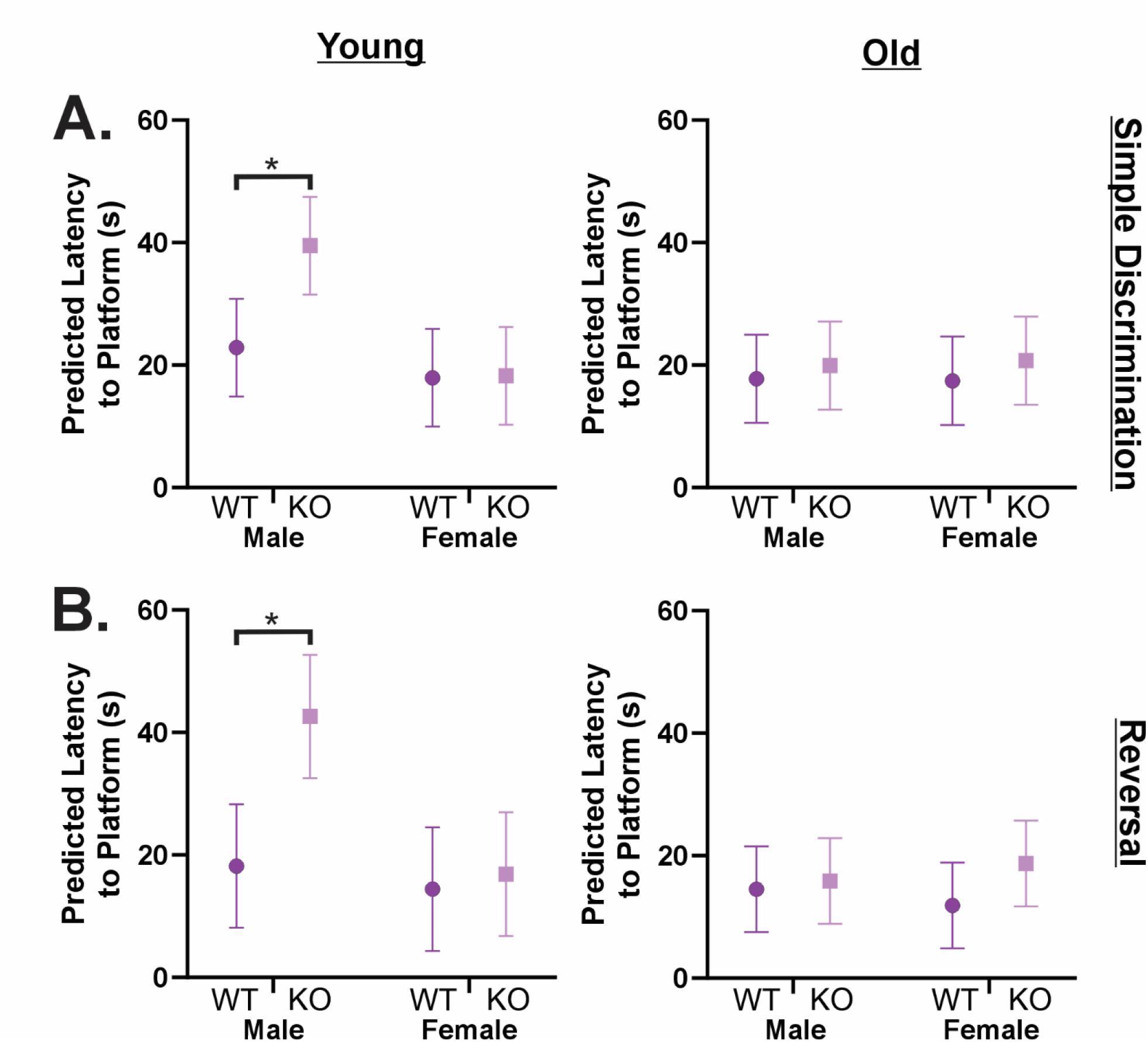
CRES KO males—but not females—exhibit deficits in spatial learning and behavioral flexibility. Learning, memory, and executive functioning were assessed in young and old WT and CRES KO mice using a two-choice discrimination water T-maze paradigm. Young and old cohorts were analyzed separately but are plotted side-by-side for ease of visualization. **A)** Simple discrimination. Statistical parameters for young mice: Genotype [*F* (1, 45.408) = 4.590, *p* = 0.038], Sex [F (1, 45.508) = 10.899, *p* = 0.002], and the Genotype x Sex interaction [F (1, 45.408) = 4.245, *p* = 0.045]. Parameters for old mice: Genotype [F (1, 50.115) = 0.570, *p* = 0.454], Sex [F (1, 50.115) = 0.004, p = 0.949], the Genotype x Sex interaction [F (1, 50.115) = 0.026, p = 0.874]. In young mice, the Genotype × Sex interaction was a significant predictor of learning and memory ability. Bonferroni-corrected post hoc comparisons demonstrated that young CRES KO males took, on average, significantly longer [16.658 s] to locate the escape platform than WT males. Older mice of either genotype did not significantly differ. **B)** Reversal learning. Statistical parameters for young mice: Genotype [F (1, 33.294) = 7.337, p = 0.011], Sex [*F* (1, 33.294) = 8.846, *p* = 0.005], and Genotype × Sex interaction [*F* (1, 33.294) = 4.908, *p* = 0.034]. Parameters for old mice: Genotype [*F* (1, 41.948) = 1.392, *p* = 0.245], Sex [*F* (1, 41.948) = 0.001, *p* = 0.982], and Genotype × Sex interaction [*F* (1, 41.948) = 0.623, *p* = 0.434]. In young mice, the Genotype × Sex interaction was a significant predictor of executive functioning and behavioral flexibility. Bonferroni-corrected post hoc comparisons demonstrated that young CRES KO males took, on average, significantly longer [19.691 s] to locate the platform than WT males. In contrast, Genotype, Sex, and their interaction were not significant predictors of cognitive performance in older mice. Data are presented as predicted values ± 95% CI.

In contrast, in aged animals, there were no significant differences between genotypes (WT, KO), sexes (male, females), or the interaction between genotype and sex, in the ability to find the escape platform. This suggests a recovery of the aforementioned cognitive deficits observed in younger mice: *Simple Discrimination*; Genotype [*F* (1, 50.115) = 0.570, *p* = 0.454]; Sex [*F* (1, 50.115) = 0.004, *p* = 0.949]; Genotype x Sex [*F* (1, 50.115) = 0.026, *p* = 0.874]; *Reversal*; Genotype [*F* (1, 41.948) = 1.392, *p* = 0.245]; Sex [*F* (1, 41.948) = 0.001, *p* = 0.982]; Genotype by Sex [*F* (1, 41.948) = 0.623, *p* = 0.434] (Fig 6A-B).

To confirm that the increased latency observed in young CRES KO males was driven by true spatial learning deficits rather than decision hesitancy or altered exploratory strategy, choice accuracy was analyzed across sequential trial blocks. These error data closely aligned with the latency measures. In young mice, a three-way repeated measures ANOVA revealed a significant Block × Genotype × Sex interaction during simple discrimination [*F* (2, 72) = 3.576, *p* = 0.033]. Probing this interaction using simple main effects demonstrated that young CRES KO males made significantly more incorrect arm entries across learning blocks compared to WT males [*p* < 0.002]. Conversely, spatial choice accuracy did not significantly differ between young WT and CRES KO females [*p* > 0.05].

This spatial deficit persisted during the reversal learning stage, indicative of impaired executive function and behavioral flexibility. Analysis of the reversal trial blocks again revealed a significant Block × Genotype × Sex interaction [*F* (2, 72) = 3.248, p = 0.045]. Consistent with the initial acquisition phase, simple main effects confirmed that young CRES KO males committed significantly more incorrect spatial choices across the reversal blocks compared to WT males [*p* < 0.001], while young CRES KO females exhibited no such deficit relative to WT controls [*p* > 0.05].

Consistent with the latency data, choice accuracy in aged mice did not significantly differ between genotypes or sexes, nor did it yield a significant three-way interaction [Simple Discrimination: *F* (2, 72) = 2.536, *p* = 0.086; Reversal: F (2, 72) = 0.091, *p* = 0.913] (Fig 7A-B). Together, this trial-block learning curve confirms that the delayed performance observed in young CRES KO males is driven by incorrect spatial choices, definitively identifying a deficit in spatial learning and executive function rather than generalized decision hesitancy.

**Figure 7.**
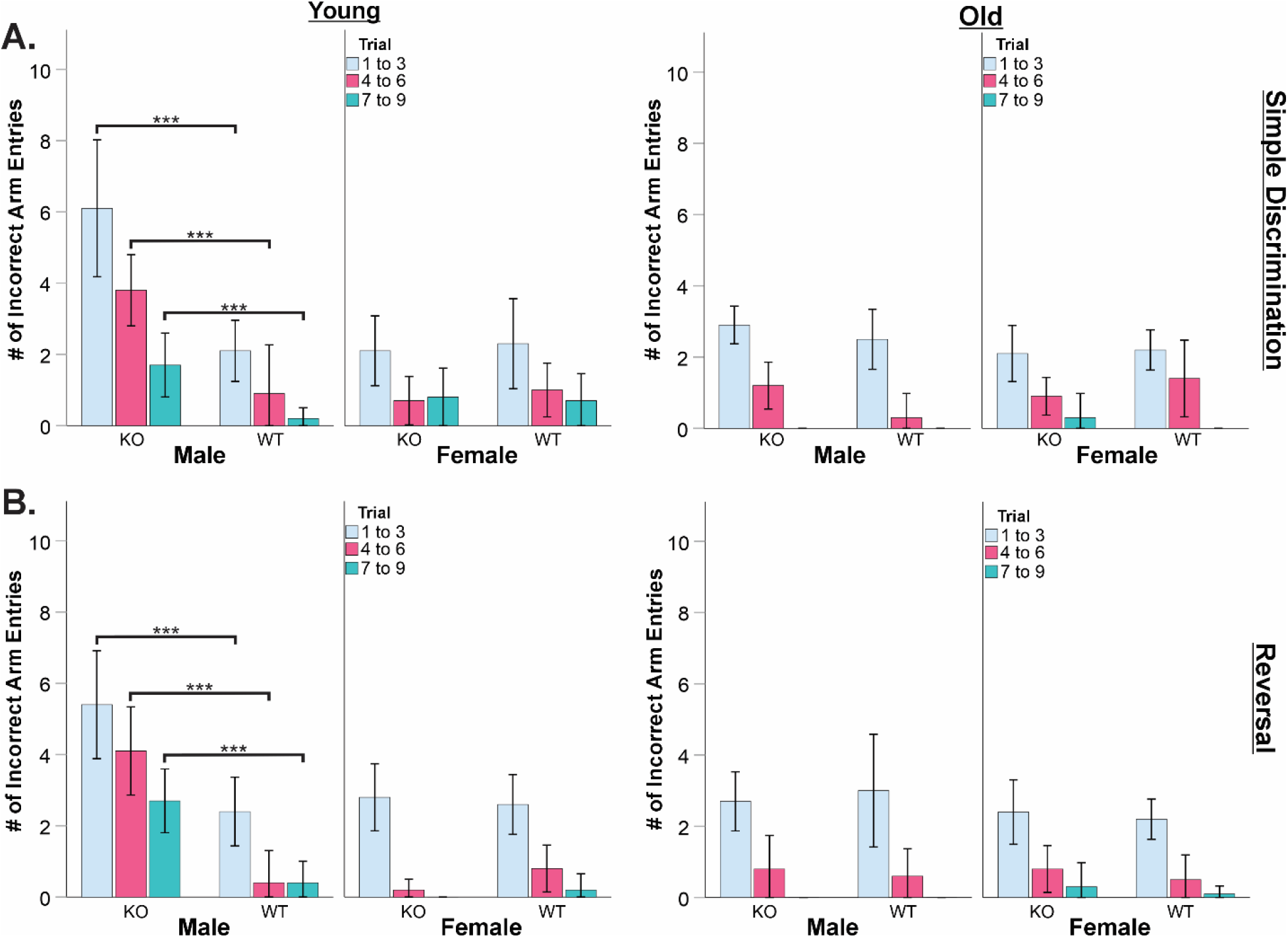
Choice accuracy confirms that delayed performance in young CRES KO males is driven by true spatial learning deficits rather than decision hesitancy. To distinguish cognitive impairment from altered exploratory strategy, choice accuracy (cumulative number of incorrect arm entries) was analyzed across sequential 3-trial blocks in young and old WT and CRES KO mice. Young and old cohorts were analyzed separately but are plotted side-by-side for ease of visualization. A) Simple discrimination. In young mice, the interaction between Block, Genotype, and Sex was a significant predictor of choice accuracy [*F* (2, 72) = 3.576, *p* = 0.033]. Probing this interaction with simple main effects demonstrated that young CRES KO males made significantly more incorrect arm entries across learning blocks compared to WT males (p < 0.002). Conversely, young CRES KO females exhibited no significant deficit relative to WT controls (p > 0.05). Older mice of either genotype or sex did not significantly differ in their spatial choice accuracy; Block × Genotype × Sex interaction [*F* (2, 72) = 2.536, *p* = 0.086]. B) Reversal learning. Consistent with the initial acquisition phase, the three-way interaction was a significant predictor of executive functioning and behavioral flexibility [*F* (2, 72) = 3.248, *p* = 0.045]. Simple main effects confirmed that young CRES KO males committed significantly more incorrect spatial choices across the reversal blocks compared to WT males (p < 0.001), while young CRES KO females did not significantly differ from WT females (p > 0.05). The interaction between Block, Genotype, and Sex was not a significant predictor of choice accuracy in older mice; Block × Genotype × Sex interaction [*F* (2, 72) = 0.091, *p* = 0.913]. Data are presented as means ± 95% CI.

Representative videos for all groups can be found on figshare (https://doi.org/10.6084/m9.figshare.31855621). These results highlight that CRES may play an age- and sex-related function in learning, memory, and executive functioning, with males lacking CRES showing a relative deficit in cognitive (learning and memory) and behavioral flexibility (executive functioning) during the early adulthood periods.

No differences were observed in voluntary locomotor behavior in the open-field apparatus and motor coordination using the accelerating rotarod in young and aged WT and KO males and females (Fig SI4). These results suggest that the lack of any observed differences in the older mice in our tests of learning, memory and executive functioning is unlikely to be solely explained by age-related impairments in motor behavior.

## Discussion

Using a CRES KO mouse model, we show here that mice lacking CRES exhibit sex-specific differences in ECM structure and brain functioning, suggesting a role for CRES as a regulatory or structural component of the brain ECM and important for learning and memory and executive functioning.

### Structural changes in loose/interstitial and PNN ECM populations in CRES KO mice

The ECM is known to be structurally different between males and females particularly in the cortex, where the male ECM has been defined as being fluid compared to the more “solid” /stiff ECM in the female (Batzdorf et al. 2022). Indeed, the composition of ECM structural proteins including collagens, laminins and fibronectin differ between the male and female cortex, resulting in distinct ECM organization and viscoelasticity, affecting brain function (Batzdorf et al. 2022). Our studies using the CRES KO model support and extend these sex-specific distinctions by showing divergent changes in hippocampal loose/interstitial ECM structure with the loss of CRES. The loose/interstitial ECM in the male KO hippocampus displayed small matrix fragments suggesting improper/incomplete assembly (as evidenced by TEM images suggestive of early amyloids) and/or possibly an overall less stable structure while in the KO female larger aggregates/assemblies were observed, indicative of a more aggregation prone and insoluble/stiff ECM. Thus, in addition to the levels of collagen and laminins contributing to differences in ECM structure between males and females, amyloidogenic/insoluble proteins such as CRES may add to the sex-specific differences in ECM stiffness/solubility.

Intriguingly, the highly branched matrix with associated spheres that is characteristic of the hippocampal WT loose/interstitial ECM is comparable in structure with the CRES subgroup-containing epididymal amyloid ECM that functions in the epididymal lumen (Do et al. 2019; Myers, Hastert, and Cornwall 2022). The similar morphologies, including the presence of sphere-like structures, between the epididymal and brain ECM implies commonalities in ECM components, such as CRES, and possibly overlapping functions. While we have yet to determine the identity of the sphere-like structures, we speculate they represent biological condensates in different states of phase separation/assembly.

We also observed changes in the structure of the condensed PNNs in male and female CRES KO hippocampus and cortex. Young male and female mice had reduced WFA staining in PNNs yet similar numbers of ACAN+ PNNs indicating that while PNNs were still present in the KO mice they were in a less mature/altered state compared to PNN structure in WT mice. Reduced WFA staining is often an indication of decreased CS chains and specifically 4-sulfation (Miyata et al. 2012). However, our sulfation analyses revealed no significant changes in the sulfation patterning, total sulfation or abundance of CSO, CSA, or CSC isomers in the CRES KO mice. While we cannot rule out the possibility that the KO genotype affected the ability of the CS chains to be extracted from the biochemical fractions, our data suggest that the loss of CRES primarily affects the CS-GAG content. It is also possible that other ECM structural proteins, independent of the CSPG aggrecan, are altered in the KO PNNs. Our studies showed that the hippocampus in young male mice was particularly affected by the loss of CRES and exhibited reduced WFA intensity and a reduced number of WFA+ PNNs in addition to a reduced number of ACAN+ PNNs. These data show that in males the number of PNNs may be reduced from an early age. However, the WFA intensity and number of ACAN+ PNNs recovered in older male mice suggesting either compensatory mechanisms or those PNNs that were present were delayed in their development. The loss of CRES also resulted in sex-specific effects in the young KO females where the number of ACAN+ PNNs in the parietal and occipital cortex were dramatically reduced but little to no change was observed in the same regions of the male KO cortex. The effects were transient since the number of ACAN+ PNNS recovered to WT levels in the older female KO mice. Our results show that CRES performs region- and sex-specific PNN functions.

### Sex-specific behavioral differences in CRES KO mice

PNNs have been shown to play a crucial role in ECM plasticity and remodeling mechanisms, and cell-to-cell communication in the brain (Van’t Spijker and Kwok 2017). Our studies show CRES is an active participant in maintaining the PNN, and loose/interstitial ECM structure, and may modulate plasticity in a region- and sex-specific manner. The PNNs in the hippocampus of young male mice were especially susceptible to the loss of CRES which may explain why males, but not females, displayed deficits in learning, memory and executive functioning.

Crucially, while young male KO mice took significantly longer to locate the escape platform, our choice accuracy data confirm this delay was driven by elevated spatial errors rather than generalized decision hesitancy or motor impairment. Young KO males not only struggled to learn the initial spatial relationship (simple discrimination), but displayed even more profound behavioral inflexibility when required to unlearn and switch to a new platform location (reversal learning). These results definitively highlight that KO males lack the cognitive flexibility required to adapt when a previously learned rule is changed. These findings are consistent with the recent work of others, where chemical manipulation of PNNs in younger mice was shown to similarly impact reversal learning in a sex-specific manner (Lyle and Verpeut 2025). Although reduced WFA intensity of PNNs is indicative of increased plasticity (Lensjo et al. 2017), in this context, it could also reflect a pathological reorganization of the ECM or a dysfunctional, heightened plasticity contributing to the observed cognitive-behavioral deficits. Importantly, the spatial learning and flexibility deficits observed in younger male KO mice were absent in older male KO mice. This behavioral rescue correlated directly with a recovery of PNN structure and maturity, and possibly a return to “normal” PNN function, as WFA intensity levels were no longer different in older KO males compared to their WT counterparts.

A decrease in WFA intensity of PNNs was also observed in young female KO mice throughout the hippocampus and cortex, but with no apparent effect on brain function based on the behavioral outcomes measured. Similarly, profound decreases in the number of ACAN+ PNNs in the parietal and occipital cortex of young KO females, as well as increased WFA intensity and a higher number of PNNs compared to WT in older females, did not translate into deleterious effects during our cognitive-behavioral evaluations. While it is possible that brain deficits not captured by our specific T-maze paradigm may occur in the KO females, the resistance of female brains to biochemical alteration without displaying overt behavioral changes has been previously reported (Malone et al. 2025). This resilience is thought to represent an evolutionary adaptation to minimize the detrimental effects of the dynamic ECM remodeling that occurs in response to fluctuating hormone levels during their rapid (4–5 day) estrous cycle. PNNs are highly sensitive to sex hormones and undergo significant reorganization in the presence of brain-derived estradiol (Ciccarelli et al. 2021; Nguyen et al. 2025; Hernández-Vivanco et al. 2022). In the results presented here, behavioral and biochemical studies of CRES were conducted in female mice without regard to the specific stage of their estrous cycle. Further studies are needed to determine if the loss of CRES in females may be more impactful behaviorally at distinct estrous stages, especially given that CRES expression in the pituitary gland is regulated by GnRH and steroid hormones (Sutton-Walsh, Whelly, and Cornwall 2006).

### Compensatory responses and/or delayed maturation in CRES KO mice

Our studies show that most of the changes in PNN structure in the young KO mice were no longer present in older KO mice. While an increase in WFA fluorescence intensity has been reported to occur normally in aged mice and is associated with an increase in inhibitory mechanisms (Bavelier et al. 2010), the older KO mice may use upregulation of sugar content as compensation for the loss of CRES. Other CRES subgroup members could also compensate for the loss of CRES in the older mice. In addition to CRES, CRES2, CRES3, and cystatin E2, which are also amyloidogenic *in vitro* and *in vivo,* are expressed throughout the brain but at varying levels between region and sex (Gomez et al. 2026). While their presence in the ECM has yet to be established, their different amyloid propensities (Whelly et al. 2016) and varying levels could further influence the “solubility”/stiffness properties of ECM throughout the WT male and female brain and differentially compensate for the absence of CRES in the KO brain. It is also possible that the development of the brain ECM is delayed in CRES KO mice resulting in loose/interstitial ECM structures that are inappropriately assembled and PNNs lacking mature CSPG content and structure. However, with age these deficits are minimized or lost as the brain ECM gradually recovers structurally and functionally.

## Conclusion

Emerging evidence suggests alterations in the brain ECM, and in particular PNN structure and function contribute to a variety of pathologies including neurodegenerative diseases and psychiatric disorders, several of which are sexually dimorphic (Sorg et al. 2016). Therefore, elucidating ECM structure/function relationships, including those that differ between males and females, are essential to understand sexually dimorphic behavior and disease. Our studies demonstrate that CRES plays a regulatory and/or structural role in the brain ECM and contributes to sex differences in ECM organization that may underlie neuroplasticity and remodeling processes that drive sex-specific behaviors. Indeed, the loss of CRES resulted in a male-specific behavioral deficit in learning and memory. Further studies are needed to determine the full extent by which ECM structure and behavior are altered in CRES KO animals including the role of other PNN components and the contributions of other CRES subgroup members.

## Acknowledgements

The authors gratefully acknowledge Mary Catherine Hastert for her assistance with the TEM imaging.

## Supplemental information

**Figure SI1.**
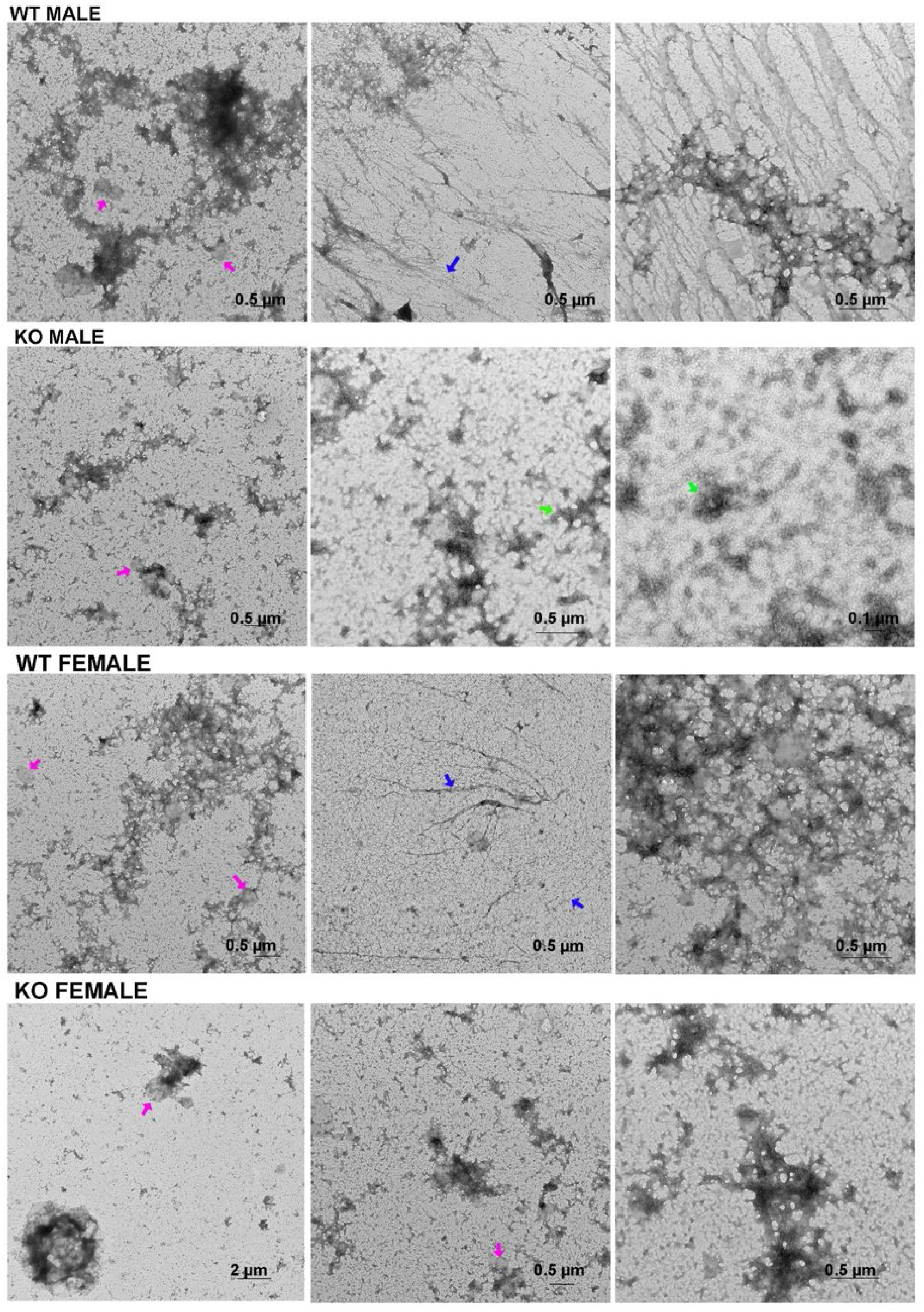
Negative stain TEM of hippocampal loose/interstitial (E1) pellets from young male and female WT and CRES KO mice revealed sex and genotype-specific differences. Scale bar, as indicated. Pink arrows, ball-like structures; blue arrows, fibrils/ thin film; green arrows, protofibril/oligomers.

**Figure SI2.**
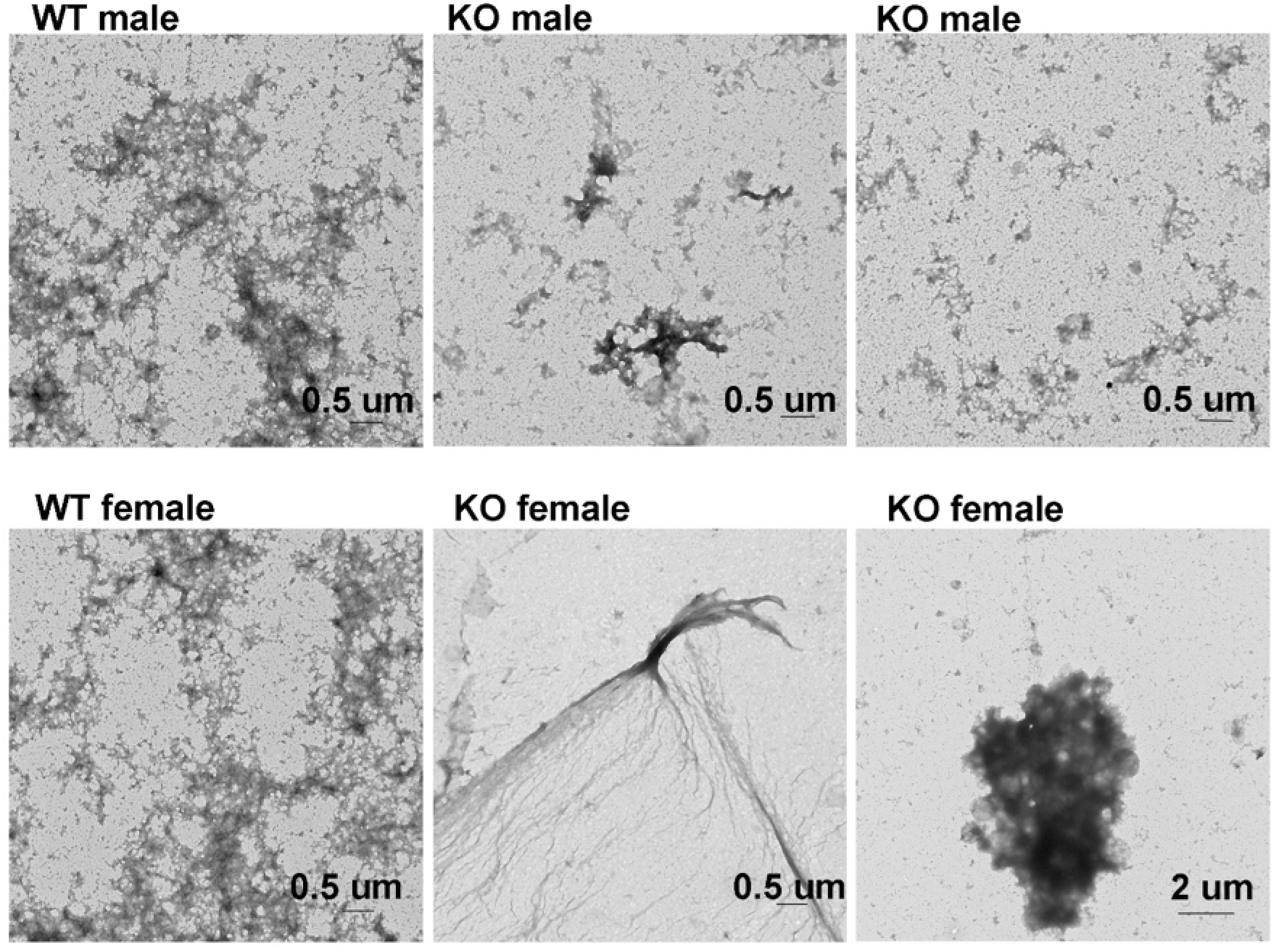
Negative stain TEM of total (not centrifuged) hippocampal loose/interstitial extract (E1) from male and female WT and CRES KO mice. Scale bar, as indicated.

**Figure SI3.**
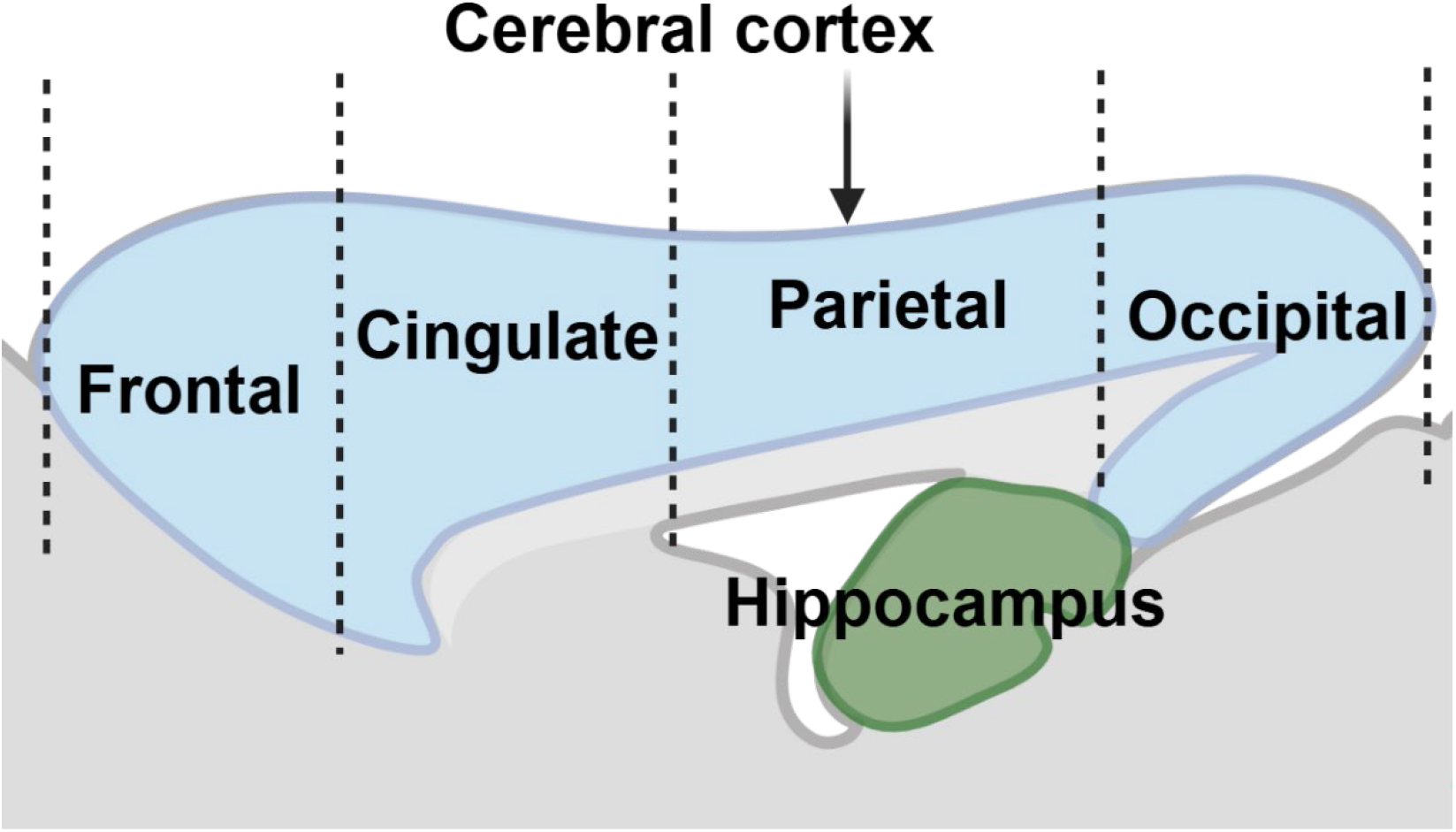
Schematic diagram showing how the cortex in sagittal sections of the mouse brain was divided into different regions for PNN analysis.

**Figure SI4.**
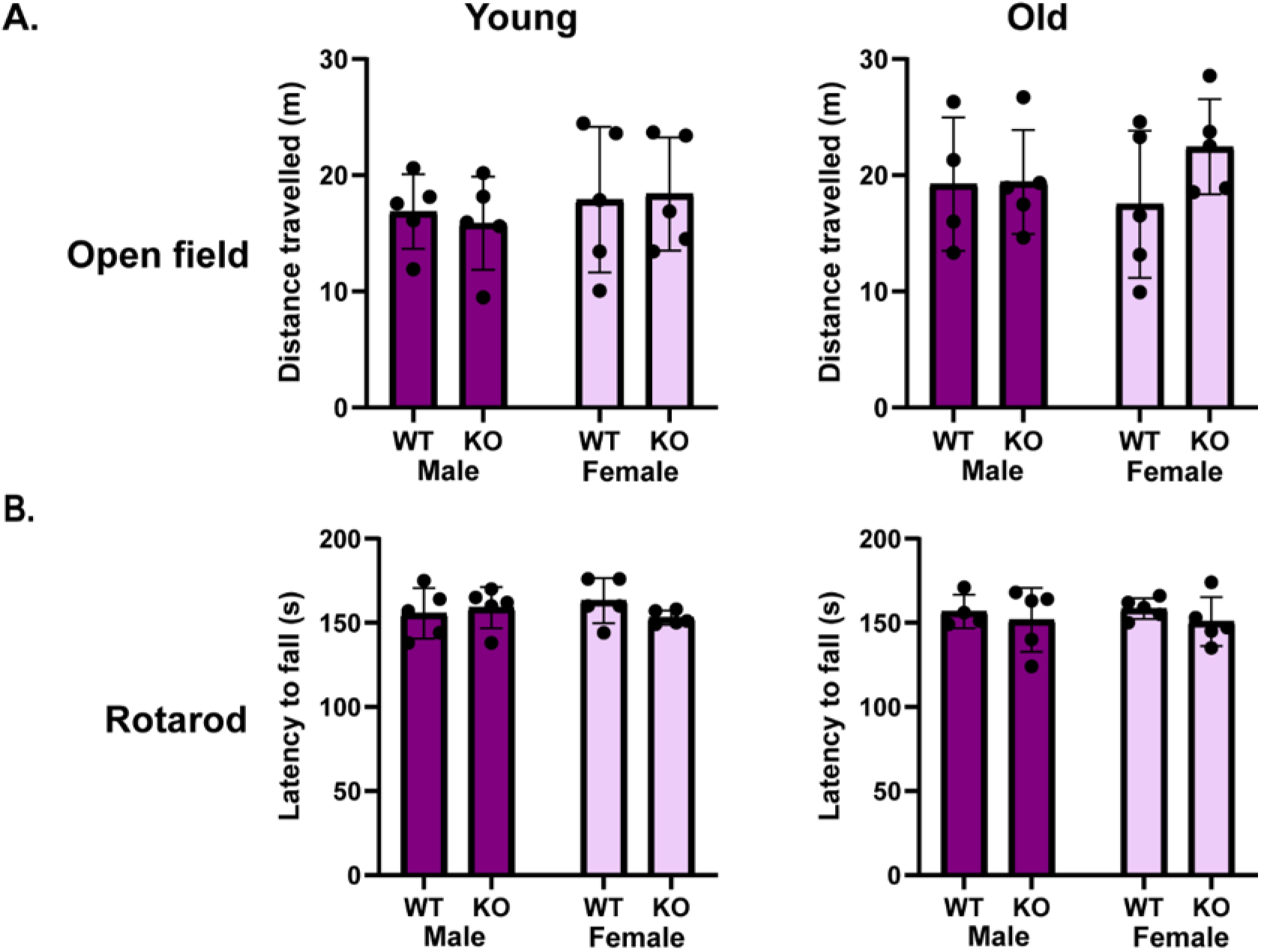
Neither young nor old CRES KO mice exhibited differences in locomotor behavior compared to WT mice, in a test of voluntary locomotion (open-field) or involuntary locomotion (the accelerating rotarod). <u>Rotarod</u> *Young mice*; genotype (*F*_1,20_ = 1.066, p = 0.317); sex (*F*_1,20_ = 0.434, p = 0.519); genotype by sex (*F*_1,20_ = 2.673, p = 0.122); *Old mice*; genotype (*F*_1,19_ = 0.070, p = 0.795); sex (*F*_1,19_ = 0.593, p = 0.453); genotype by sex (*F*_1,19_ = 3.341, p = 0.088); <u>Open-Field,</u> *Young mice*; genotype (*F*_1,20_ = 4.124, p = 0.059); sex (*F*_1,20_ = 0.134, p = 0.719); genotype by sex (*F*_1,20_ = 1.614, p = 0.222); *Old mice*; genotype (*F*_1,19_ = 0.145, p = 0.709); sex (*F*_1,19_ = 0.006, p = 0.940); genotype by sex (*F*_1,19_ = 1.835, p = 0.196). Values represent Mean ± SD of n= 5 mice except for one old WT male mouse that was excluded due to technical failure.

